# Reduction of auditory input improves performance on the heartbeat tracking task, but does not necessarily enhance interoception

**DOI:** 10.1101/830141

**Authors:** Jennifer Todd, Farah Hina, Jane E. Aspell

**Affiliations:** School of Psychology and Sport Science, Anglia Ruskin University, Cambridge, UK

## Abstract

Previous research utilising a between-subjects design has indicated that the use of noise-dampening ear-protectors might enhance interoceptive accuracy (IAcc). In the present study, we further examined this effect using a repeated-measures, within-participants design, and investigated potential mechanisms that might explain the effect. 50 participants completed the heartbeat tracking task (HTT) with and without the use of industrial ear-protectors, in a counter-balanced order. Participants were asked to count the number of heartbeats occurring in five discrete time intervals of 25, 35, 45, 55 and 95 seconds, without feeling for a manual pulse. HTT scores were significantly higher when ear-protectors were worn, and the improvement in performance was greatest for participants with lower baseline IAcc. The ear-protectors were associated with significantly increased self-reported heartbeat audibility, task-related confidence and concentration, and decreased levels of distractibility. Heartbeat audibility was also correlated with HTT performance when the ear-protectors were worn. Because the use of industrial ear defenders resulted in increased heartbeat audibility, this manipulation should not be used to assess causal hypotheses related to changes in IAcc. However, it may serve as a simple, non-invasive manipulation to assess the effects of ‘externalised’ interoceptive signals.

**Highlights:**

- Ear-protectors elicit an improvement in performance on the heartbeat tracking task
- This improvement is greatest for participants with lower baseline accuracy
- The improvement is associated with the auditory perception of the heartbeat
- Prior use of ear-protectors did not result in later enhanced performance on the control condition without ear-protectors.

---

*Interoception* refers to the brain’s processing of internal physiological stimuli: information regarding the present condition of the body (e.g. heart rate, temperature, blood sugar levels) is detected, interpreted and integrated within the nervous system to provide a continual indication of the body’s internal state, across both conscious and subconscious levels (Khalsa et al., 2018). Interoceptive signalling is a crucial component of homeostatic functioning (Barrett, & Simmons, 2015), and has also been associated with aspects of higher order cognition, such as decision making (e.g., Dunn et al., 2010; Piech et al., 2017; Werner et al., 2013), and emotion perception and regulation (e.g., Herbert, Pollatos & Schandry, 2007; Kever, Pollatos, Vermeulen, & Grynberg, 2015; Pollatos, Matthias, & Keller, 2015). In addition, people with certain clinical conditions may have levels of interoceptive accuracy that are higher or lower than the general population (Khalsa et al., 2018; Quadt, Critchley & Garfinkel, 2018).

Research supports that interoception is a multi-faceted construct (Forkmann et al., 2016; Garfinkel, Seth, Barrett, Suzuki & Critchley, 2015; Khalsa et al., 2018), and scholars typically distinguish between three facets: *interoceptive accuracy* (IAcc), which describes the objective detection and tracking of internal bodily sensations with behavioural measures such as heartbeat perception tasks (e.g. Schandry, 1981; Whitehead, Drescher, Heiman & Blackwell, 1977); *interoceptive sensibility* (IS) which refers to self-reported awareness of internal bodily sensations via questionnaires (e.g. Mehling et al., 2012), or perceived task performance (Garfinkel et al., 2015); and finally *interoceptive awareness* (IAw), which refers to the ‘metacognitive’ correspondence between IAcc and confidence ratings (Garfinkel et al., 2015).

Investigations of IAcc have been dominated by assessments of cardiac perception accuracy, often to the exclusion of other bodily domains (Khalsa et al., 2018). Indeed, it is frequently implied that cardiac perception accuracy signifies a marker of overall interoceptive ability (Tsakiris & Critchley, 2016). Whilst some literature supports this notion (Herbert, Muth, Pollatos & Herbert, 2012), it has not yet been fully explored (Khalsa et al., 2018). Moreover, the reliability of the heartbeat tracking task (HTT; Schandry, 1981) – the most commonly used measure of IAcc – has been heavily challenged recently. The task may be confounded by participants’ abilities to estimate time (Ainley, Brass, & Tsakiris, 2014; c.f. Shah, Hall, Catmur & Bird, 2016), participants’ knowledge/expectations about their resting heart rate (Murphy, Millgate, et al., 2018; Knapp-Kline & Kline, 2005; Ring, Brener, Knapp & Mailloux, 2015), and affected by physiological factors such as age, body mass index (BMI), blood pressure, resting heart rate, and heart rate variability (Khalsa, Rudrauf, & Tranel, 2009; Murphy, Geary, Millgate, Catmur, & Bird, 2018; Zamariola, Maurage, Luminet, & Corneille, 2018). Furthermore, a number of problems have been identified regarding HTT scores. For example, while HTT scores should theoretically be unbiased to error type (i.e., the over- or underestimation of heartbeats should be equally weighted), Zamariola and colleagues (2018) found that over 95% of scores reflect underestimation of heartbeats, and that the overestimation of heartbeats is disproportionately associated with higher HTT scores. Moreover, the specific wording of the task instructions (Desmedt, Luminet & Corneille, 2018), the length of the time intervals used for the task (Zamariola et al., 2018; cf. Tsakiris, Ainley, Pollatos, & Herbert, 2019), and the equipment utilised to measure heartbeats (Murphy et al., 2019) can all have significant effects on task performance. Therefore, further research scrutinising and improving the task methodology is needed.

Whilst the available interoception literature predominantly focuses upon group differences in trait IAcc and IS, (e.g Ateş Çöl, Sonmez & Vardar, 2016; Klabunde, Acheson, Boutelle, Matthews & Kaye, 2013; Pollatos et al., 2008), there is also evidence to suggest that interoceptive processing can be manipulated as a state variable, over both short and long-term periods. For example, short-term manipulations of IAcc include mirror self-observation (Ainley, Tajadura Jiménez, Fotopoulou & Tsakiris, 2012), focusing upon self-referential information (Ainley, Maister, Brokfeld, Farmer & Tsakiris, 2013), and performance-related feedback in a cardiac discrimination task (Piech et al. 2017). Meanwhile, long-term manipulations of IS and IAcc include meditation and mindfulness interventions (Bornemann, Herbert, Mehling & Singer, 2015; Fischer, Messner & Pollatos, 2017; Kok & Singer, 2017; c.f. Khalsa, Rudrauf, Hassanpour, Davidson, & Tranel, 2019; Parkin et al., 2014). Another potential manipulation of cardiac perception accuracy may involve reducing external auditory input. Indeed, in a recent quasi-experimental study, Hall, Lopes and Yu (2019) demonstrated that the use of noise-dampening ear-protectors resulted in significantly higher scores on the HTT (Schandry, 1981). However, due to the utilisation of a between-subjects study design and given the large variability in heartbeat detection scores (Murphy, Brewer, Hobson, Catmur, & Bird, 2018), it is not possible to ascertain from the work of Hall and colleagues (2019) whether IAcc can be manipulated within individuals by use of ear protectors. Indeed, whilst the difference in IAcc reported by Hall and colleagues (2019) may be a direct result of the ear-protectors, it could also be a reflection of pre-existing group differences in IAcc, or even factors that have associated with cardiac perception accuracy such as body BMI, heart rate variability, blood pressure, or knowledge of resting heartrate (Murphy, Geary, et al., 2018; Murphy, Millgate et al., 2018), particularly as allocation to conditions was not randomised. It was also unclear from the work of Hall and colleagues (2019) *why* the use of ear-protectors might increase interoceptive accuracy. One hypothesis is that the ear-protectors reduce background auditory noise, making the heartbeat more discriminable. Another possibility is that the protectors act as a resonant medium, audibly amplifying the heartbeat pulse or making it perceptible via tactile sensations caused by pressure on a blood vessel.

The primary aim of the present study was to assess whether the recent findings from Hall and colleagues (2019) could be replicated utilising a repeated-measures design, which would control for the possible confounding effects of induvial differences previously outlined. We expected that the use of noise-dampening ear-protectors would elicit a significant improvement in performance on the HTT. Secondarily, we sought to explore potential mechanisms that might explain differences in performance across the two conditions, by asking participants to self-report: (1) levels of perceived task performance (task-related confidence); (2) the degree to which they could concentrate on the task; (3) the degree to which they felt they could ‘hear’ their heartbeat (‘heartbeat audibility’); and (4), the degree to which they felt distracted during the task. Previous research has identified negative relationships between cardiac IAcc and age (Murphy, Geary, et al., 2018), BMI (Herbert, Blechert, Hautzinger, Matthias, & Herbert, 2013; Herbert & Pollatos, 2014; Murphy, Geary et al., 2018) and resting heartrate (Zamariola et al., 2018), therefore, additional effects of these variables were also anticipated.

## Method

### 1. Participants

An *a priori* power analysis indicated that a sample of 27 participants would be sufficient to detect a medium-sized effect of the ear protectors upon HTT-scores, at α = .05, with power at .80. To detect a medium-sized effect of the ear protectors upon the four VAS indices, 43 participants would be sufficient to achieve power at .80, with α = .0125. In practice, a sample of 50 staff and students were recruited from Anglia Ruskin University. The sample was comprised of 21 men and 28 women, and one person who described their gender as ‘other’. Participants were aged between 18 and 42 (*M* = 26.08 ± 6.73), and BMI values ranged from 16.30 to 34.80 kg/m^2^ (*M* = 24.24 ± 4.91). The distribution of the data for BMI is comparable to the most recent data on British adults (National Health Service Digital, 2016)^1^.

Participants were ineligible if they had any known auditory deficits, neurological/psychological conditions, or cardiovascular conditions. Participation was voluntary, and participants were not offered any form of remuneration.

### 2. Materials and Apparatus

A PowerLab device (AD Instruments, Oxford, Oxfordshire, UK) was connected to a PC to record participants’ electrocardiograms (ECG) and hence their heartbeats throughout the experiment; data was recorded using LabChart 8 software. Three disposable electrocardiography (ECG) electrodes (positioned on the chest in standard three-lead configuration), relayed R-wave output through shielded wires. The electrodes were self-attached by participants under their clothes, guided by a visual diagram.

A pair of 3M Peltor Optime adjustable ear-protectors, with a single number rating (SNR) noise reduction of 27 dB (recommended for moderate industrial noise), were used during the experimental condition.

### 3. Design

All participants were asked to complete the HTT (Schandry, 1981 see section 4) under two conditions: during the experimental condition, participants wore the ear-protectors described in section 2.2, and during the control condition participants completed the task without the ear-protectors. The conditions were completed in two separate blocks in a counterbalanced order: participants randomly allocated even numbers completed the experimental condition prior to the control condition, whilst participants who were allocated odd numbers completed the control condition prior to the experimental condition.

### 4. Procedure

The study was approved by the authors’ university ethics committee prior to data collection. All testing took place in a university laboratory, with only the researcher and the participant present. There were no obvious visual or auditory distractions present during testing sessions and the windows within the laboratory remained shut. The foyer area outside of the laboratory was also vacant during all testing sessions. Participants were first presented with a written information sheet, and then asked to provide written informed consent.

Throughout the HTT, participants were seated in an upright position facing away from the experimenter. Participants were asked to attempt to sense their heart beating in the inside of their body, without using a manual pulse or watch. Consistent with the recommendation of Desmedt and colleagues (2018), participants were asked to count and report only the number of heartbeats actually felt, without attempts to guess heart rate. Participants were asked to count the number of heartbeats occurring in five discrete time intervals of 25, 35, 45, 55 and 95 seconds (presented in a random order across participants). The researcher delivered vocal ‘start’ and ‘stop’ cues which indicated the beginning and end of each trial. Prior to the commencement of the experimental condition, it was ascertained that the participants were comfortable wearing the ear-protectors, and that they could accurately hear and respond to the auditory start and stop signals. Participants were unaware of how long they were counting for, and no performance-related feedback was given.

Immediately following the completion of each HTT block (experimental/control), participants were asked to respond to the following questions using a computer which was situated adjacent to the experimental set-up:

1. How confident are you in your responses for the last five trials?
2. To what extent did you feel you could concentrate during the last five trials?
3. To what extent did you feel you could ‘hear’ your heartbeat during the last five trials?
4. To what extent did you feel you were distracted by background noises during the last five trials?

The questions were presented in a random order across participants. Responses were recorded on a visual analogue scale (VAS) that was 10cm long which participants could mark using a computer mouse. The left end was marked “Not at all” and the right end was marked “Completely”. Participants were also offered the opportunity to write any additional comments about the tasks. Finally, participants were asked to complete a demographic questionnaire. At the end of the study, all participants were presented with written debriefing information.

## Results

### 5.1 Transformation of raw data

Interoceptive accuracy scores from the HTT were calculated using the following equation: IAcc = (1 – [recorded heartbeats – counted heartbeats]/recorded heartbeats), as in Schandry (1981). Scores were calculated for each trial, and then averaged across five trials to generate an overall interoceptive accuracy score for each condition. Absolute values were utilised, so that scores range from 0 to 1 and those closer to 1 indicate greater IAcc.

Meanwhile, each VAS score ranged from 0 and 100, with scores incrementing at a rate of 1 point per mm, such that positioning the cursor at the far left of the scale (“Not at all”) was assigned a score of 0, and positioning the cursor at the far right of the scale (“Completely”) was assigned a score of 100 (as in Pfeifer et al., 2017).

### 5.2 Interoceptive Accuracy

A paired samples *t*-test was used to determine whether there was a statistically significant difference in interoceptive accuracy when the HTT was completed with and without the use of ear-protectors. Inspection of the difference scores for the experimental and control conditions revealed no outliers (as assessed by a boxplot), and the scores were normally distributed (as assessed by Shapiro-Wilk’s test, *p* = .706). As can be seen from Figure 1, participants tended to demonstrate greater interoceptive accuracy when using ear-protectors (*M* = .86 ± .12), compared to without (*M* = .59 ± .20), a statistically significant difference in heartbeat perception scores of .26 (95% CI, .204 to .325), *t*(49) = 8.764, *p* = < .0005, *d* = 1.24.

**Figure 1.**
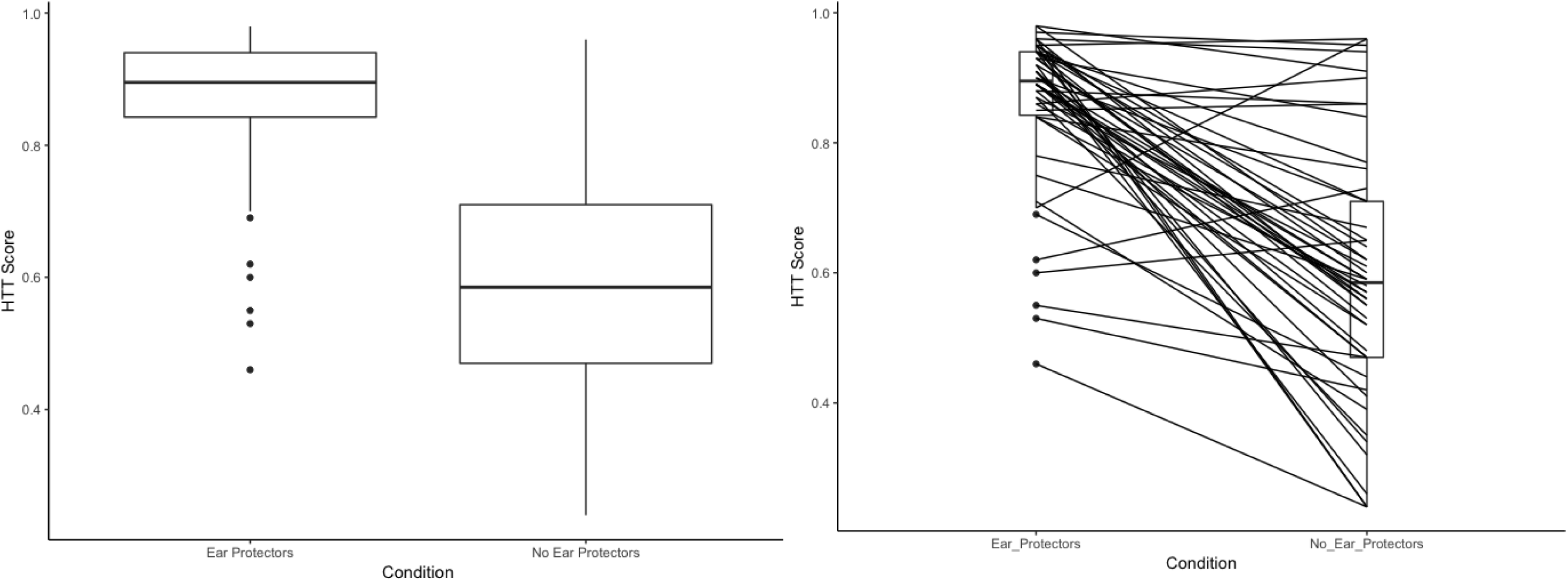
Box plots showing (left) the distribution of scores on the heartbeat tracking task across the two conditions, and (right) changes in the performance for individual participants across the two conditions.

Next, we sought to determine whether there was an effect of order (completion of the control/experimental task first/second) upon task scores. To control for the multiple comparisons, a Bonferroni correction was applied, such that *p* = .05/2= .025. For the control condition, inspection of boxplots revealed no outliers, and the data were normally distributed for participants who completed the condition first and second (Shapiro-Wilk’s test: *p* = .657, .168, for each group). An independent-samples *t-*test revealed that cores for the control condition were slightly higher for participants who completed it before the use of the ear-protectors (*M* = .60 ± .17) compared to those who completed it after (*M* = .58 ± .23), but the mean difference of .020 (95% CI, .094 to .134) was not statistically significant, *t*(48) = - 1.112, *p* = .725, *d* = .099. For the experimental condition, the data were negatively skewed (the z-scores for skewness were -3.55 and -3.91, and Shapiro-Wilk’s test: *p* < .001 for both groups), therefore a Mann-Whitney U test was used. Median interoceptive accuracy scores did not significantly differ between groups *U* = 392, *z* = 1.543, *p* = .123.

In order to assess whether scores for the ear protectors condition differed according to baseline performance on the control condition, the HTT performance data were split into two groups by the median (as in Ainley et al., 2012; 2013), such that participants who scored ≤ .58 were classified as ‘low’ cardiac perceivers, and participants who scored ≥ .59 were classified as ‘high’ cardiac perceivers (for each group, *n* = 25). We then conducted two tests to examine (1) whether ‘low’ and ‘high’ cardiac perceivers differed in performance during the experimental condition; and (2) whether the difference scores (i.e. performance on the control condition, subtracted from performance when the task was completed with ear protectors) varied across ‘low’ and ‘high’ perceivers. An adjusted significance threshold was applied: *p* = .05/2= .025. The data for the ear protectors condition were not normally distributed for ‘low’ or ‘high’ perceivers (Shapiro-Wilk’s test: *p* ≤ .001 for each group), therefore a Mann Whitney-U test was utilised. There was no statistically significant difference between the two groups, *U* = 365.00, *z* = 1.019, *p* = .308. The difference scores were normally distributed (Shapiro-Wilk’s test: *p* = .706) and inspection of boxplots indicated that there were no outliers. An independent-samples *t-*test revealed an effect of baseline sensitivity, such that participants who were classified as ‘low’ cardiac perceivers had a greater mean improvement (.40), than participants who were classified as ‘high’ cardiac perceivers (.13), a statistically significant mean difference of .27, (95% CI, .1.79 to 3.66), *t*(48) = 5.86, *p* = < .0005, *d* = 1.66.

### 5.3 Task reliability

Cronbach’s *α* was .85 for the control condition, and .92 for the experimental condition. Friedman tests were run to determine if task performance differed by trial length (25, 35, 45, 55 and 95 seconds) for each condition. There were no statistically significant differences for either the control condition, *χ*^2^(4) = 1.081, *p* = .897, or the experimental condition, *χ*^2^(4) = 1.896, *p* = .755.

### 5.4 Visual Analogue Scales

Wilcoxon signed-rank tests were used to assess the self-reported data from the VASs. As can be seen from Table 1, the use of ear-protectors elicited significant changes in four aspects of HTT experience after accounting Bonferroni correction (*p* = .05/4 = .0125): heartbeat audibility, HTT-related confidence, HTT-related concentration, and HTT-related distractibility.

**Table 1.** The number of participants reporting an altered heartbeat tracking task experience as a result of the noise dampening ear protectors, and results of Wilcoxon signed-rank tests.

| Aspect of HTT experience | Reported<br>Increase (*n*) | Reported<br>Decrease (*n*) | No<br>change<br>(*n*) | Control<br>condition<br>median<br>rating | Experimental<br>condition<br>median<br>rating | Median<br>change | *z* | *p* |
| --- | --- | --- | --- | --- | --- | --- | --- | --- |
| Performance-related<br>confidence ratings | 30 | 7 | 3 | 50.00 | 72.50 | 16.00 | 5.16 | < .0005 |
| Task-related concentration<br>ratings | 39 | 11 | 0 | 60.00 | 80.00 | 21.50 | 4.67 | < .0005 |
| Heartbeat audibility | 44 | 3 | 3 | 27.50 | 80.00 | 30.00 | 5.36 | < .0005 |
| Distraction | 6 | 32 | 12 | 20.00 | 0.00 | -12.00 | 4.31 | < .0005 |

*Notes.* *N* = 50, HTT = Heartbeat Tracking Task.

Next, Spearman’s correlations were conducted to ascertain the relationship between performance on the HTT, the self-reported VAS variables, and additional demographic factors (see Table 2). Given the large number of comparisons, we controlled for false discovery rate (FDR) using the Benjamini and Hochberg (1995) procedure. Additionally, differences in the pattern of the correlation coefficients across the experimental and control conditions were compared by computing Hittner, May, and Silver’s (2003) modification of Dunn and Clark’s (1969) *z*, using the cocor package (Diedenhofen, & Musch, 2015) in *R* (Rosseel, 2012). Performances across the two conditions were not strongly correlated, and below the threshold for statistical significance. Of all the VAS variables, heartbeat audibility had the strongest correlation with task performance during the experimental condition (*r_s_ =*

**Table 2.** Correlations between heartbeat tracking task scores, heartbeat tracking task performance-judgements, and demographic variables within the experimental condition (top diagonal) and within the control condition (bottom diagonal).

|  | HTT score | Confidence | Concentration | Audibility | Distracted | Age | BMI | Resting heartrate |
| --- | --- | --- | --- | --- | --- | --- | --- | --- |
| HTT score | **.138** | **.400\*\*** | **.342\*** | **.456\*\*** | **-.225** | **.405\*\*** | **-.103** | **-.433\*\*** |
| Confidence | .389** | **.538\*\*** | **.720\*\*** | **.798\*\*** | **-.366\*\*** | **.071** | **-.062** | **-.324\*** |
| Concentration | .347* | .825** | **.418\*\*** | **.605\*\*** | **-.504\*\*** | **-.134** | **-.122** | **-.109** |
| Audibility | .178 | .533** | .458** | **.346\*** | **-.246** | **.019** | **-.208** | **-.237** |
| Distracted | .050 | -.185 | -.307* | -.210 | **.323\*** | **.112** | **-.175** | **.092** |
| Age | -.049 | -.105 | -.062 | -.317* | <.001 | - | **.114** | **-.491\*\*** |
| BMI | .079 | -.187 | .137 | .205 | -.211 | .114 | - | **-.059** |
| Resting Heartrate | .035 | -.065 | -.130 | .178 | .150 | -.491** | -.059 | - |

*Notes.* $N = 50$ ; \* $p$ -FDR < .05; \*\* $p$ -FDR < .01. HTT = heartbeat tracking task; BMI = Body mass index. Correlations between variables within the experimental condition (top diagonal), are presented in bold for clarity. Correlations for the same variable across the two conditions (diagonal line) are also presented in bold for clarity.

.456; see Table 2). However, it was not significantly associated with IAcc within the control condition. Regarding the remaining VAS variables, there were significant, positive associations between performance-related confidence ratings and HTT scores, and between concentration ratings and HTT scores, and these associations did not differ significantly across the two conditions (Table 3). Conversely, self-reported distraction ratings were not significantly associated with IAcc on either condition (Table 3). In terms of the inter-correlations between the VAS variables, there was a strong, significant (*r_s_ = .*798; *p* > .001) correlation between confidence and audibility within the experimental condition. Heartbeat audibility and concentration ratings were significantly associated (*r_s_ = .*605; *p* > .001) within the experimental condition. Finally, age was significantly, positively correlated with IAcc, and negatively correlated with resting heartrate within the experimental condition, but the variables were not significantly associated within the control condition.

**Table 3.**
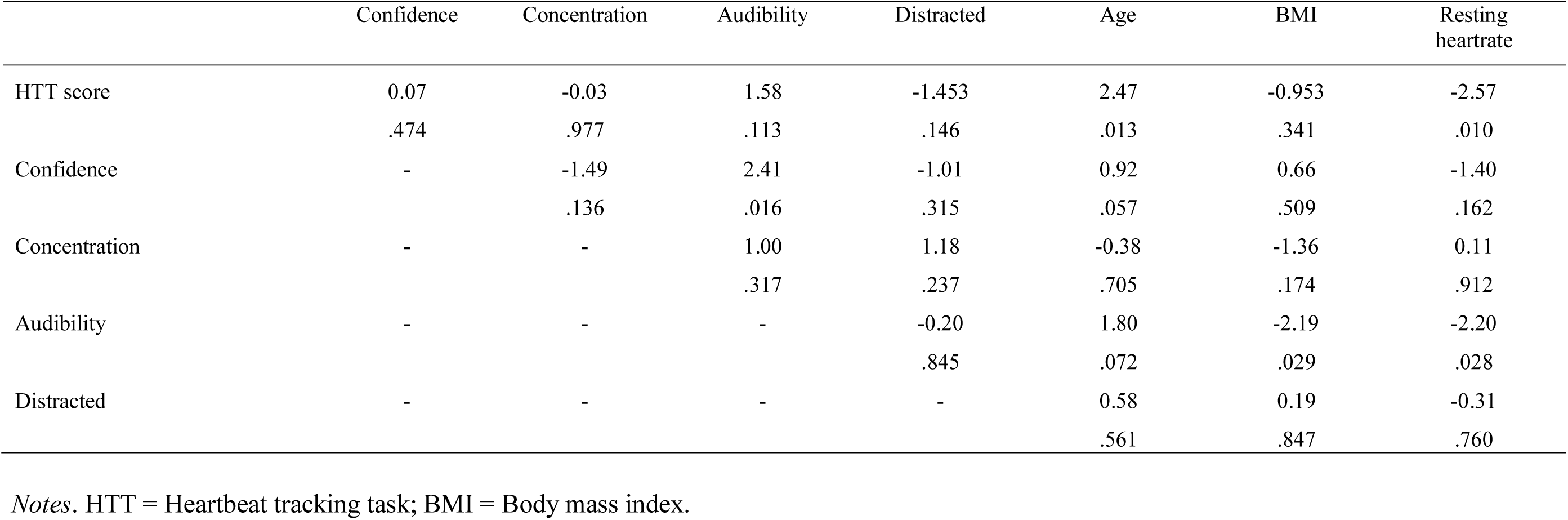
z_observed_ values and associated p values for comparison of the correlation coefficients across the experimental and control conditions.

|  | Confidence | Concentration | Audibility | Distracted | Age | BMI | Resting<br>heartrate |
| --- | --- | --- | --- | --- | --- | --- | --- |
| HTT score | 0.07<br>.474 | -0.03<br>.977 | 1.58<br>.113 | -1.453<br>.146 | 2.47<br>.013 | -0.953<br>.341 | -2.57<br>.010 |
| Confidence | - | -1.49<br>.136 | 2.41<br>.016 | -1.01<br>.315 | 0.92<br>.057 | 0.66<br>.509 | -1.40<br>.162 |
| Concentration | - | - | 1.00<br>.317 | 1.18<br>.237 | -0.38<br>.705 | -1.36<br>.174 | 0.11<br>.912 |
| Audibility | - | - | - | -0.20<br>.845 | 1.80<br>.072 | -2.19<br>.029 | -2.20<br>.028 |
| Distracted | - | - | - | - | 0.58<br>.561 | 0.19<br>.847 | -0.31<br>.760 |

Notes. HTT = Heartbeat tracking task; BMI = Body mass index.

## Discussion

The aim of the present research was to assess whether the recent findings from Hall and colleagues (2019) - regarding the effects of ear protectors on IAcc - could be replicated using a repeated-measures, within-participant design. In accordance with our hypothesis, there was a significant increase in heartbeat tracking task (HTT) scores when the task was performed using ear-protectors compared to without. In addition, the improvements in performance were significantly greater for participants who had lower scores on the control condition. We also sought to assess possible mechanisms which might explain this effect. In doing so, we found that the use of ear-protectors was associated with participants’ ratings of heartbeat audibility; confidence in perceived task performance; the degree to which participants felt they could concentrate; and, the degree to which participants felt distracted throughout the task.

Performances across the two conditions were not significantly correlated (*r_s_* = .138, *p* = .338), suggesting that participants who performed well in one condition did not necessarily perform well in the other. Indeed, our data show that participants who had lower scores on the control condition exhibited significantly greater increases in HTT-scores in the experimental condition than participants who had higher scores on the control condition. Conversely, participants with ‘low’/’high’ scores on the control condition did not differ significantly their performance in the experimental condition. That is, the use of the ear protectors appears to have somewhat homogenised performance on the HTT. These findings may reflect that the two conditions are best considered as different tasks performed using different cues.

There are several possible mechanisms that might explain the distinction between the two conditions. One explanation centres upon the finding that participants reported the heartbeat pulse to be more audible when wearing the ear-protectors. The ear protectors are designed to fit closely to the head, and it might be that the pressure of the ear-protectors caused perceptible sensations via pressure on a blood vessel. As one participant reported after wearing the ear-protectors, “it [my heartbeat] was easier to hear in this condition, but the beat was in my head, not my heart.” It is possible that the ear-protectors alter the body location in which the heartbeat is experienced (it is usually experienced in the chest region, and sometimes the neck; Nummenmaa, Hari, Hietanen, & Glerean, 2018), although this supposition requires further investigation. Nevertheless, the current evidence suggests that the ear-protector manipulation transformed the HTT into a task that is inherently exteroceptive: HTT scores were significantly higher when ear-defenders were worn, and participants reported that the heartbeat pulse was more audible while wearing the ear-defenders. Therefore, whilst scores on the HTT tend to improve as a result of the ear-protectors, it should not be asserted that the manipulation increases IAcc (as suggested by Hall et al., 2019), given that IAcc purportedly measures sensitivity to *internal* bodily signals.

It is possible that the ear-protector HTT manipulation could be used examine the interplay between interoceptive and exteroceptive cues, or the effects of ‘externalised’ interoceptive signals. For example, previous research has identified that externalised heartbeat signals affect bodily self-consciousness (Aspell et al., 2013), chronic pain (Solca et al., 2018), and social preferences such as altruism and fairness (Lenggenhager, Azevedo, Mancini, & Aglioti, 2013). The use of ear-protectors could serve as a simple, non-invasive way to ‘externalise’ interoceptive stimuli. Indeed, within the present study some participants reported that the ear-protectors amplified other bodily processes too, for example one participant commented that “breathing and swallowing became suddenly distracting”.

Alternative explanations for the distinction between the two conditions draw upon the remaining self-report variables. While heartbeat audibility had the greatest median change across the two conditions, the use of the ear-protectors was also associated with significant changes in the following variables (in order of effect size): (1) performance-related confidence; (2) the extent to which participants felt they could concentrate during the task, and; (3) the extent to which participants felt distracted during the task. According to the competition of cues hypothesis (Pennebaker, 1982; Pennebaker & Lightner, 1980), attention is continually divided across internal and external stimuli, such that attention toward external stimuli will reduce the attentional resources available for internal stimuli (and *vice versa*). As such, the reduction of exteroceptive stimuli via the ear protectors is likely to have increased the salience of interoceptive cues, making them easier to report accurately. However, the study setting was quiet, and the conditions within the laboratory were controlled during the study. Due to the relatively small sample, it was not possible to calculate the amount of variance in HTT scores that can be attributed to each of the VAS variables. Furthermore, it is also likely that this effect is not specific to ear-protectors: the reduction of visual input through use of a blind-fold, for example, might also promote the salience of interoceptive stimuli.

It is important to note that the present work has several additional limitations. First, the ear protectors utilised in the present work had a moderate noise-reduction rating, and it is possible that the present findings may not generalise to other types of ear-protector. For example, noise-cancelling headphones with a lower noise-reduction rating might not be associated with changes in HTT scores. Should researchers choose to include ear-defenders in future HTT protocols, we recommend that the possible performance-related effects of the equipment are tested beforehand, and that the specifications of the equipment are clearly reported. Second, while the procedure was implemented by the same researcher across all participants, it is possible that the vocal delivery of the HTT instructions was may have introduced a small degree of error into the data. Future work should address this issue using a computer-based protocol. Finally, participants completed the VASs at the end of each condition which may have encouraged desirability bias in the reporting of task-related experience.

These limitations notwithstanding, the findings from the present study support the findings of Hall and colleagues (2019) utilising a within-participants repeated-measures design. Our results suggest that the associated improvement in performance on the heartbeat tracking task was partially due to self-reported increases in the audibility of the heartbeat sound. The exteroceptive nature of the feedback from the ear-protectors therefore precludes the utility of this specific type of equipment to assess causal hypotheses related to changes in IAcc. However, the ear-protector manipulation may serve as a simple, non-invasive paradigm to assess the interplay between interoceptive and exteroceptive cues, or the effects of ‘externalised’ interoceptive signals (e.g. Aspell et al., 2013; Lenggenhager et al., 2013).

## Footnotes

1 As some previous HTT research has been conducted with participants who have BMI values between 18.5 and 25, we also completed a series of analyses using a subset of participants from the present study who were within this BMI category. We can confirm that the results from these analyses did not differ from the results presented in the current manuscript.

